# A Metacell Model of Single Cell RNA-seq Counts Yields a Gaussian Mixture Model in PCA Space

**DOI:** 10.64898/2026.09.21.753263

**Authors:** Sivan Leviyang

## Abstract

Single cell RNA sequencing (scRNAseq) has revolutionized cellular biology over the past decade. At its most basic, a scRNAseq dataset is a cell by gene matrix with entries giving the transcription count of a particular gene in a particular cell. A common workflow applies log-normalization and z-scoring to the count matrix and then embeds the cells in a lower dimensional space using principal component analysis (PCA). A large number of computational tools use the PCA embedding as a starting point, but statistical models for the distribution of cells in the PCA space are lacking. As a result, many tools depend on ad-hoc assumptions that likely lead to signal loss.

In this work, we address this gap using the metacell model, a statistical model of count matrices introduced by previous authors. We show that application of the workflow to count matrices generated under a metacell model yields PCA embeddings distributed as a Gaussian mixture model (GMM). To our knowledge, this is the first result connecting a statistical model of counts to the distribution of cells in PCA space. Our derivation depends on methods from random matrix theory (RMT) and we extend and establish connections between RMT and scRNAseq count matrices.

We then use the predicted GMM to test the metacell model on 11 datasets. The predictions are accurate for counts generated by the metacell model but underestimate the variance of real cells for most of our datasets. We show that this error largely arises from gene correlations within metacells, which the metacell model does not capture. Some of our datasets are fit relatively well by the metacell model, suggesting that the fit of the metacell model to true datasets depends on sufficiently dense sampling. Finally, we show that the strength of noise in the PCA space varies significantly across metacells, suggesting that statistical modeling of PCA embeddings can improve existing downstream workflows.

Our results provide a framework for deriving and testing statistical models of PCA embeddings from statistical models of counts and identify a limitation of the metacell description for scRNAseq datasets.

---

Single cell RNA sequencing (scRNAseq) has revolutionized cellular biology over the last decade, leading to a wide array of computational challenges [57, 27, 58]. Most scRNAseq computational workflows start with a count matrix in which each row corresponds to a cell, each column to a gene, and the entries give the expression count of a gene for a cell. Typically, the count matrix is in some way normalized and then the rows, i.e. cells, are embedded in a lower dimensional space. A common workflow, implemented as the default in the Seurat package and supported by the scanpy package [61, 22], applies log-normalization and z-scoring to the count matrix followed by a principal component analysis (PCA) that embeds the cells in a lower dimensional space: the PCA space. The embedding computed by this workflow is used in a wide array of downstream analyses and algorithms [42]. For example, the scRNAseq batch correction algorithm Harmony operates on the cells in PCA space instead of the original counts [31]. And perhaps the most commonly used scRNAseq algorithms, cell clustering for the de-novo identification of cell types and UMAP dimension reduction for visualization, both construct nearest neighbor graphs based on cell distances in PCA space [30, 36, 7].

Despite the central role of this workflow - hereafter referred to as the standard workflow - in scRNAseq analyses, little is known about how signal and noise in the observed cell counts are transformed by the standard workflow into signal and noise in the PCA embedding of the cells. In particular, we lack a statistical model of cell noise in PCA space. In contrast, noise models for the count matrix - typically involving the negative binomial distribution - play a central role in count matrix pre-processing [20, 56, 59, 35, 16]. For example, the gene selection algorithms used in scanpy and Seurat apply a variance scaling based on the negative binomial distribution [55, 21]. Picking genes according to unscaled variance preferentially selects housekeeping genes, leading to loss of biological signal. Such effects may be occurring downstream of the standard workflow. If different cells have different variance levels and geometries in PCA space, the numerous algorithms that work downstream of the PCA embedding may be losing biological signal.

In this work, we address the following question: given a generative model for scRNAseq counts, what distribution is induced on cells in the PCA space by the standard workflow? A wide array of generative models have been developed for scRNAseq count matrices [52, 40, 63, 53]. We consider the metacell model, introduced and developed by previous authors [6, 8, 12, 13, 39, 49]. The definition of a metacell varies across authors, but intuitively a metacell is a collection of cells in which count variation arises due to uncorrelated noise across genes and cells and not due to a biological signal. Given a scRNAseq count matrix, we partition the cells into metacells and model expression of all cells in the metacell as negative binomial distributed. The metacell model then gives a statistical model of the dataset and we can generate a count matrix by sampling counts for each cell in each metacell.

We show that under a metacell model, cells within a metacell are distributed as a multivariate normal (MVN) in the PCA space, making the PCA embedding across metacells a Gaussian mixture model (GMM). We derive the mean and variances parameterizing the GMM as a function of the negative binomial parameters in the metacell model, providing an analytic connection between the count matrix and the cell distribution in PCA space. To our knowledge, this is the first result that connects a statistical model for cell counts to a distribution on the PCA space.

The metacell model is an approximation and it does not necessarily accurately predict the PCA embedding of true (i.e. not generated in-silico) scRNAseq datasets. We therefore ask whether PCA embeddings of true datasets are well fit by GMMs with parameters predicted by the metacell model. We find that for some datasets, the metacell model’s generated GMM distribution in PCA space is a good fit for the true PCA embeddings, but in general this is not the case. For most of the datasets we investigate, the metacell model underestimates the variance of true PCA embeddings. The failure of the metacell model can arise from gene distributions that are not well modeled by the negative binomial and from the absence of gene correlation within a metacell. We find that the absence of gene correlation leads to most of the variance underestimate.

Based on our 11 datasets, we highlight a trend suggesting that the metacell model fails when sampling is not sufficiently dense, but further work is needed using a larger number of datasets to verify this effect. We also investigate variance in the PCA space across metacells. In PCA embeddings generated under the metacell model and in true datasets, we find metacells span a roughly 10-fold range in the variance of their cell embeddings. Under our metacell model, variance within a metacell is generated solely by sampling noise, meaning that many downstream tools that use PCA distances may be confusing noise with biological signal.

Our analysis of the metacell model uses tools from random matrix theory (RMT) [11, 9]. The connection between RMT and scRNAseq datasets was first noted by Aparicio et al. [3]. Aparicio et al. did not consider the full standard workflow, but instead considered the count matrix after log-normalization and z-scoring, which we refer to as the scaled count matrix. The authors showed empirically that after removal of outlier eigenvalues, the remaining eigenvalues - the so-called bulk eigenvalues - of the scaled count matrix have a Marchenko-Pastur (MP) distribution, a well known distribution in RMT [44]. Subsequent authors have noted a deviation of the bulk eigenvalues from the MP distribution [33, 15, 54, 32].

The results of Aparicio et al. raised the intriguing possibility that scaled count matrices can be modeled and analyzed by RMT methods. Subsequent groups have exploited RMT methods to develop scRNAseq algorithms [46, 33, 15, 54, 14, 28, 45, 29, 32, 37]. For example, Stanley et al. [54] introduce the principled PCA. Roughly, they apply a transform to the scaled count matrix to force the bulk eigenvalues to satisfy a MP distribution, which they then exploit to better distinguish outlier from bulk eigenvalues, leading to a PCA embedding of the cells that preserves more biological signal. Chardes [15] applies a similar transform and uses a sparse PCA with penalty parameter chosen using RMT, again leading to an improved PCA embedding.

Our work differs from these previous works by assuming a statistical model for the count matrix, which we then transform into a statistical model for the scaled count matrix by applying the log-normalization and z-scoring steps of the standard workflow. We then use RMT methods to derive an analytic distribution for the bulk eigenvalues of the scaled count matrix and the outlier eigenvalues and eigenvectors that form the PCA embedding. In the process of deriving our results, we also show why some scRNAseq datasets are well described by an MP distribution, as noted by Aparicio et al., while others deviate from it.

## 1 Results

Let *X* be a cell by gene count matrix of dimension *C* ×*G*. Below, we will consider count matrices with 1000− 5000 genes, reflecting the common range of highly variable genes (HVG) seen in scRNAseq studies, and 2, 500 to 180, 000 cells, see Table 1. The standard workflow involves a log-normalization [43, 1],

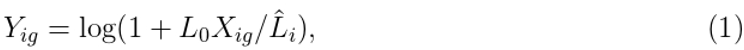

where 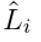is the library size of the *i*th cell, i.e. 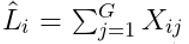 and *L*_0_ = 10000 is a common choice and one we will use. *Y* is then z-scored column-wise (i.e. gene-wise),

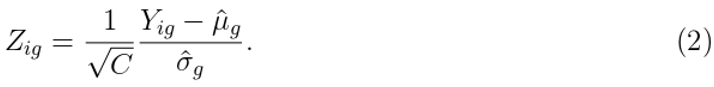

**Table 1:**
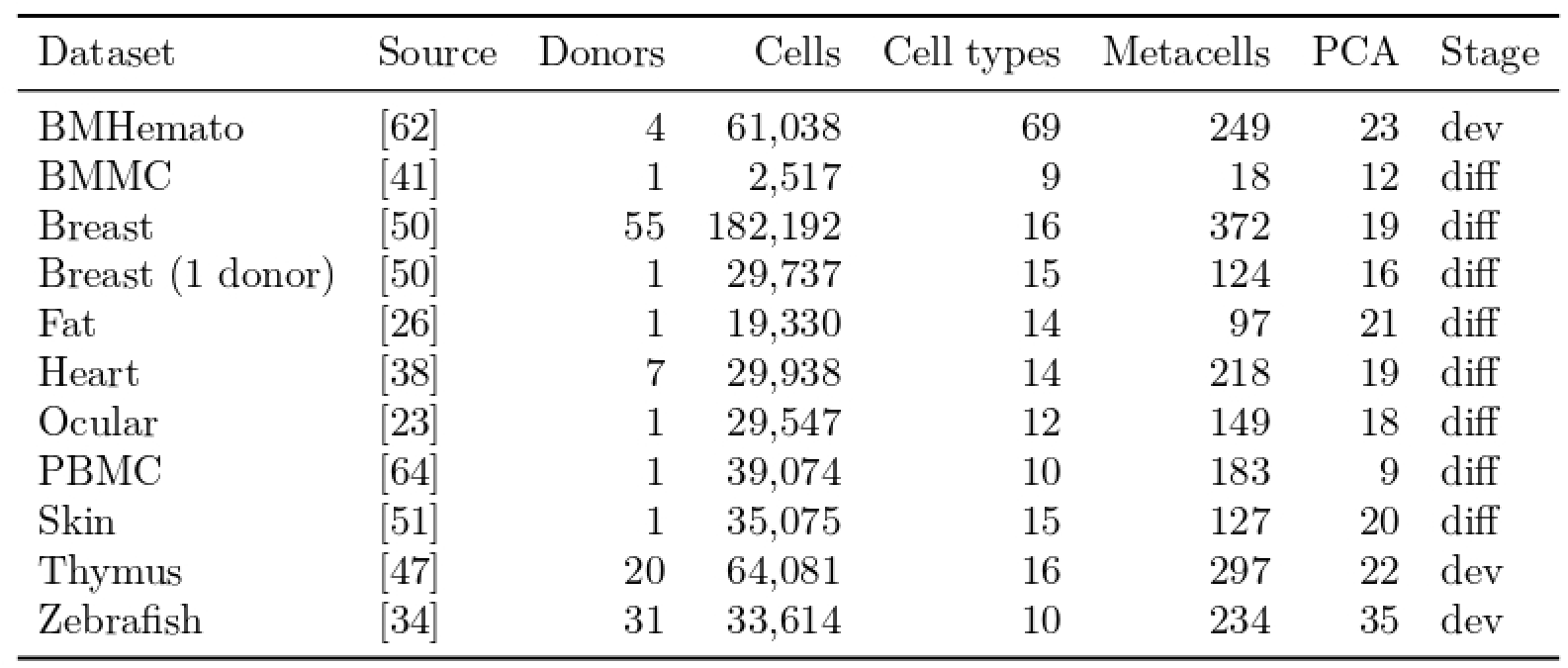
Single-cell RNA-seq datasets used in this study. All datasets were generated on the 10x Chromium platform. Donors is the number of individuals contributing cells (for Zebrafish, the number of individual embryos and larvae). PCA is the dimension *k* of the PCA space. Stage is dev (developmental) or diff (differentiated).

| Dataset | Source | Donors | Cells | Cell types | Metacells | PCA | Stage |
| --- | --- | --- | --- | --- | --- | --- | --- |
| BMHemato | [62] | 4 | 61,038 | 69 | 249 | 23 | dev |
| BMMC | [41] | 1 | 2,517 | 9 | 18 | 12 | diff |
| Breast | [50] | 55 | 182,192 | 16 | 372 | 19 | diff |
| Breast (1 donor) | [50] | 1 | 29,737 | 15 | 124 | 16 | diff |
| Fat | [26] | 1 | 19,330 | 14 | 97 | 21 | diff |
| Heart | [38] | 7 | 29,938 | 14 | 218 | 19 | diff |
| Ocular | [23] | 1 | 29,547 | 12 | 149 | 18 | diff |
| PBMC | [64] | 1 | 39,074 | 10 | 183 | 9 | diff |
| Skin | [51] | 1 | 35,075 | 15 | 127 | 20 | diff |
| Thymus | [47] | 20 | 64,081 | 16 | 297 | 22 | dev |
| Zebrafish | [34] | 31 | 33,614 | 10 | 234 | 35 | dev |

Above 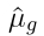 and 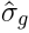 are the mean and standard deviation of the *g*th column of *Y*. We will call *Z* the scaled count matrix. (Note that *Z* and *X* have the same dimensions, *C*×*G*.)

The last step of the standard workflow is a PCA on *Z* that embeds the *C* cells into a PCA space. The dimension *k* of the PCA space is sometimes chosen as a default value. For example in Seurat *k* = 50 is the default for the RunPCA function but downstream functions such as FindNeighbors default to *k* = 10. But a more principled approach, which we take, is to choose *k* by identifying the elbow of *Z*’s eigenvalues. Letting *u*^(*i*)^ and *s*_*i*_ be the left singular vectors and singular values of *Z*, respectively, for *i* = 1, 2, …, *G* (here we assume *G* ≤ *C*, which is the case for all our datasets), the PCA embedding of *Z* is defined by the *C* × *k* dimensional matrix *W*,

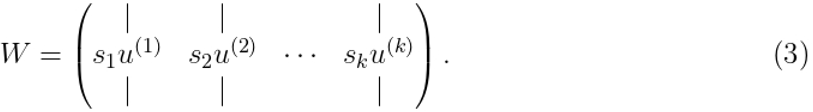

The idea of a metacell was introduced by Baran et al. [6] as a collection of cells that have the same cell count distribution up to sampling noise. Previous groups have focused on algorithms that partition cells in a dataset into metacells - see [12] for a review of the metacell concept and algorithms - but here we use it as a generative model. Recently, Liu and Li defined a metacell as a collection of cells with gene counts drawn independently from a Poisson and we take our inspiration from this definition [39]. We define the metacell model as a random count matrix *X*_metacell_ with fixed deterministic dimension *C*× *G* in which the rows (i.e. cells) are deterministically partitioned into metacells within which all cells have the same distribution. We use the negative binomial as the count distribution to allow for more flexibility than a Poisson. We assume, for cell *i* of metacell *a* and gene *g*,

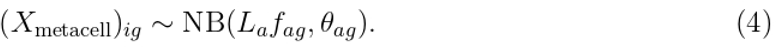

Above, NB is the negative binomial distribution, here parameterized by mean and dispersion, and each entry of *X*_metacell_ is chosen independently. The parameters *f*_*ag*_ and *θ*_*ag*_ represent the frequency and dispersion of gene *g* for metacell *a. L*_*a*_ represents a mean library size for the metacell. (We note in passing that *L*_*a*_ could be a random variable from which each cell in the metacell draws a library size; our methods would work for a random *L*_*a*_, but we have not pursued this modeling approach.) Given a scRNAseq dataset, we form a corresponding metacell model by choosing the metacell partition and fitting the negative binomial parameters of each metacell to its counts.

The difficult part of fitting a scRNAseq dataset is choosing the metacells. Once metacells are chosen, the library size *L*_*a*_ can be chosen as the mean library size of the metacell’s cells and scRNAseq algorithms exist to fit the negative binomial parameters of the metacell. We use the scTransform algorithm [21] as implemented in Seurat to infer *f*_*ag*_ and *θ*_*ag*_ once cells are partitioned into metacells. Several authors have developed algorithms that partition cells into metacells from a dataset’s count matrix [6, 8, 13, 49], but the theory is not fully developed and current metacell algorithms are not guaranteed to construct true metacells. We use the Supercell algorithm [13] to select an initial number of metacells and then add a second step in which we further refine the metacell partition, see Methods for details.

Given a metacell model, the distribution of *X*_metacell_ is determined and we can form a random scaled count matrix *Z*_metacell_ through application of the standard workflow to *X*_metacell_. Since the distribution of *X*_metacell_ is known, *Z*_metacell_ can be decomposed into a deterministic and random matrix which we call the spike and bulk matrices, respectively, following RMT terminology,

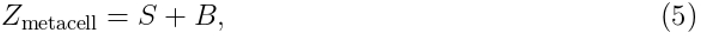

where *S* = *E*[*Z*_metacell_] and *B* = *Z*_metacell_ −*E*[*Z*_metacell_] (here *E*[] is expected value). *S* is a deterministic matrix which we can compute from the negative binomial parameters of the metacells. We let *r* be the rank of *S* and *m* the number of metacells; we always have *r* = *m*− 1. (The minus one comes from the centering of *Y*.) *B* is a mean-zero matrix with independent (but not identically) distributed entries for which we can compute the entry wise variance, see the Methods for details on these computations. Importantly, all rows of *S* in the same metacell are identical and carry the biological signal. The matrix *B* captures all of the stochasticity in the negative binomial and represents noise. *B* has the form of a mixture model with the metacells as the mixtures: each cell in a metacell draws its counts according to the metacell’s count distribution.

From an RMT perspective, *Z*_metacell_ is an example of a spiked random matrix [24, 4, 5, 48, 10]. Spiked random matrices have received extensive attention in the RMT literature, see [25] for a review. The work of Benaych-Georges and Nadakuditi [11] provides explicit formulas for the outlier eigenvalues and eigenvectors of a spiked matrix given the spike matrix *S* and the eigenvalues of the bulk matrix *B*. The work of Benaych-Georges and Couillet [9] computes the eigenvalues for random matrices in which the rows are drawn from a mixture model, exactly the case for our bulk matrix *B*.

However, these two previous works each have assumptions that are not met by *Z*_metacell_. Benaych-Georges and Nadakuditi assume that *B* is isotropic relative to *S* which is not true in our case. *S* is formed from the means of negative binomials while *B* is formed from the same negative binomials after centering. Since the variance of a negative binomial is a function of its mean, *S* and *B* are not isotropic relative to each other. Benaych-Georges and Couillet assume random matrices with normally distributed entries, while the entries of *B* are centered negative binomials. Besides the specific assumptions of these two RMT works, the accuracy of the results depends on *B* being in a universality regime, meaning that its eigenvalues and eigenvectors depend only on *B*’s variance structure - not on the particular distribution of its entries - and are accurately described by a large matrix limit in which *C, G* → ∞ with the ratio *C/G* fixed [2]. Whether *B* as specified by the metacell model is in a universality regime is a priori unclear.

To address these issues, we define a random matrix with normally distributed noise,

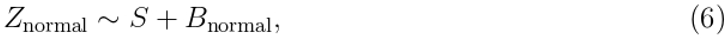

where *B*_normal_ is a mean-zero random matrix with each entry normally distributed with variance equal to the variance of the corresponding entry in *B. B*_normal_ satisfies the assumptions of Benaych-Georges and Couillet and we derive the associated large matrix limits for its eigenvalues and eigenvectors. Note that *S* and *B*_normal_ are still not isotropic. To apply the results of Benaych-Georges and Nadakuditi, we extend their results to the case of non-isotropic *S* and *B*, see Methods for details.

We show in the Methods that in the large matrix limit, the metacells of *Z*_normal_ have an MVN distribution in PCA space and we compute the mean and variance of each metacell from *S* and the variances of *B*. We refer to these results as the LML-RMT (large matrix limit, random matrix theory) predictions for the PCA embeddings.

Most of the rest of this paper evaluates the accuracy of LML-RMT predictions. Given a scRNAseq dataset, let 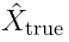 be the dataset’s count matrix. Here the hat emphasizes that 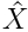is not random and the *true* subscript emphasizes that the counts are from a scRNAseq dataset and are not generated. We apply the standard workflow to 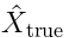 to form the scaled count matrix 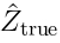. We fit a metacell model to 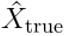, sample from the model to form 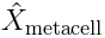and then apply the standard workflow to form a scaled count matrix 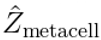.

For each dataset we construct the (non-random) matrices 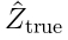and 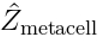. For both of these, we numerically compute eigenvalues (i.e. the square of the singular values) and left singular vectors. Given the metacell model, we can also determine the distribution of *Z*_normal_ and the spike-bulk matrices *S* and *B*_normal_. Using *S* and the variances of *B*, we can derive the LML-RMT predictions for the singular values and vectors that form the PCA embedding. Below, we evaluate the accuracy of the LML-RMT predictions relative to the singular values and vectors of 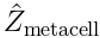 and 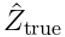.

We use 11 scRNAseq datasets described in Table 1. The datasets span a range in terms of the number of cells sampled, the number of donors, the number of cell types and whether the dataset captures a developmental trajectory of cell types or cell types from differentiated tissues.

Before proceeding to describe our experiments, we provide some qualitative orientation using UMAPs. Here, the UMAPs are intended for orientation only; below we provide rigorous justification of the fits. Figure 1A shows UMAPs of the PCA embeddings given by the LML-RMT predictions and those computed using the metacell model generated count matrix and the true count matrix for the PBMC and Thymus datasets. For the PBMC dataset, all the UMAPs are similar, reflecting a good fit of the LML-RMT to both the metacell model and true counts. For the Thymus dataset, the LML-RMT and metacell model UMAPs are similar, but the true UMAP differs: in the true UMAP, cell types have more variance.

**Figure 1:**
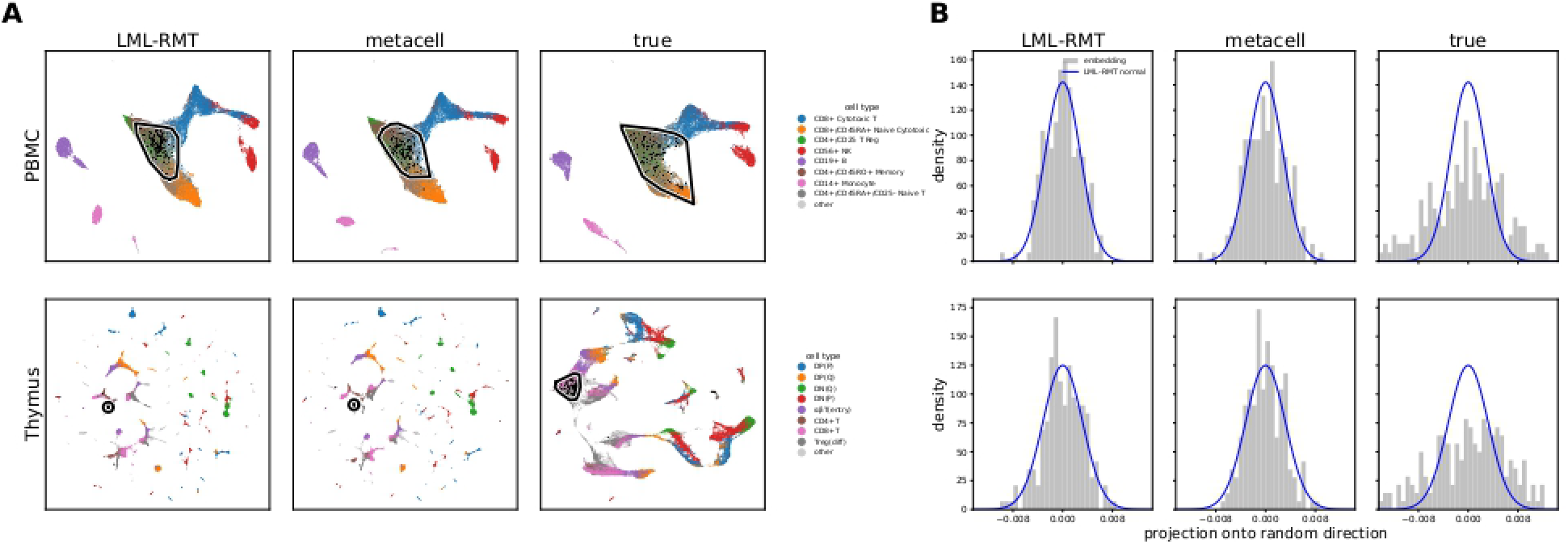
Qualitative Comparison of the LML-RMT, Metacell Model, and True PCA Embeddings. (A) UMAPs of the PCA embeddings for the PBMC (top row) and Thymus (bottom row) datasets. Each column shows the UMAP generated from the PCA embedding formed from (left) the LML-RMT prediction, (center) the metacell model scaled count matrix, and (right) the true scaled count matrix. The black outline in each panel encloses the cells of the metacell used in (B). (B) For each dataset, a single metacell with roughly 200 cells was chosen. In each PCA embedding, the cells of the metacell were centered by subtracting the metacell’s mean, and then projected onto a single random direction in PCA space, with the same direction used for all three embeddings. The grey histograms show the distribution of the projected values for the LML-RMT prediction (left), the metacell model (center), and the true count matrix (right). The blue curve in each panel is the normal distribution predicted by LML-RMT along the chosen direction.

Figure 1B shows an example of the distribution of cells in the PCA space (not in the UMAP space). For each of the PBMC and Thymus datasets, we arbitrarily chose a metacell with roughly 200 cells. Each of these roughly 200 cells has an embedding in the PCA space, which is 9 and 22 dimensional for the PBMC and Thymus datasets, respectively. We selected a random direction in PCA space and projected the cell embeddings onto that direction. The figure shows a histogram of the projection for the LML-RMT predictions and the metacell model and true PCA embeddings. The metacell model PCA embeddings are well fit by the LML-RMT predictions while the true embeddings are not. Further, the variance of the LML-RMT embeddings underestimates the variance of the true embeddings. The underestimate connects back to the UMAP in which the 200 cell metacell we selected is isolated in the LML-RMT and metacell model UMAPs for the Thymus dataset, but is connected to other metacells in the true UMAP. In the true embedding, the variance of the metacell leads to its overlap with other metacells, while the underestimate of the LML-RMT prediction leads to isolated metacells in the PCA space.

We now turn to a more rigorous evaluation of the LML-RMT predictions. Below we will refer to the eigenvalues and eigenvectors of the scaled count matrix *Z*, by which we mean the eigenvalues of *Z*^*T*^ *Z* (or equivalently the squared singular values of *Z*) and the left singular vectors of *Z*, respectively.

### 1.1 Bulk and Outlier Eigenvalues

The eigenvalues of scRNAseq scaled count matrices split into two groups: the bulk eigen-values, a tightly spaced collection of many eigenvalues, and outlier eigenvalues, a few widely spaced eigenvalues lying above the bulk eigenvalues. In the case of the spike-bulk decomposition (6), the bulk eigenvalues correspond to those of *B*_normal_ while the outliers arise from the interaction between *S* and *B*_normal_. The PCA embedding is formed from the outlier eigenvalues and their eigenvectors (recall here we mean the left singular vectors), so it is the outliers that ultimately concern us. We nevertheless begin with the bulk eigenvalues for two reasons. First, the bulk eigenvalues have been the focus of previous RMT work on scRNAseq. Second, the bulk distribution is central to the LML-RMT predictions as described in the Methods. As a tangential benefit to investigating the bulk eigenvalues, our results explain the seeming discrepancy in the literature between the Marchenko-Pastur (MP) fit found by Aparicio et al. and the deviations found by others.

Figure 2 shows the bulk eigenvalue distribution of the LML-RMT predictions against the bulk eigenvalues of 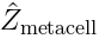and 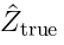 for the case of 1000 genes (i.e. *G* = 1000, see Methods for details on gene selection) for four of the datasets. Fits to the other seven datasets were similar. For 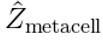and 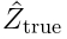we form the bulk eigenvalues by computing all the eigenvalues and dropping the outliers. The bulk eigenvalues of the LML-RMT predictions are constructed analytically as described in the Methods. The MP distribution is also shown and is determined solely by the dimensions *C* ×*G*. For brevity, we refer to these eigenvalues as LML-RMT, metacell, true, and MP eigenvalues, respectively. The metacell eigenvalues are well approximated by the LML-RMT eigenvalues. The true eigenvalues have some deviation from the LML-RMT eigenvalues, but the LML-RMT eigenvalues are overall a much better approximation than the MP distribution.

**Figure 2:**
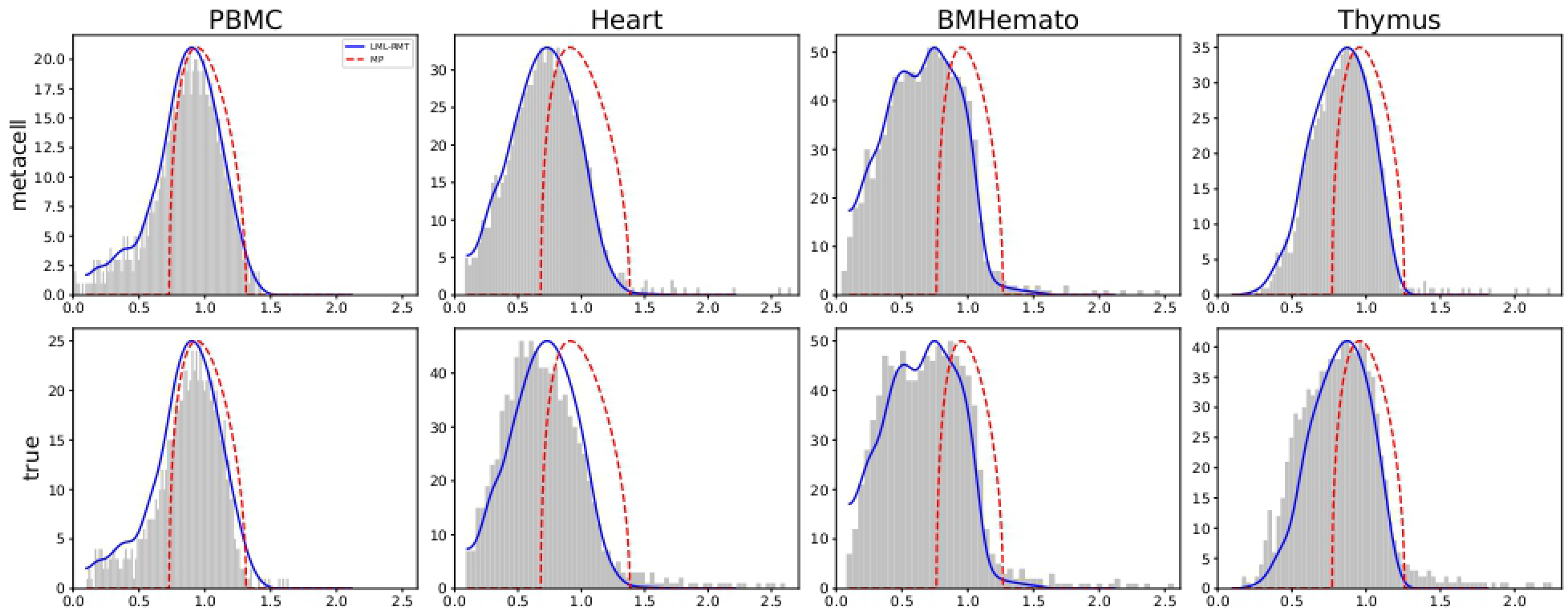
Bulk Eigenvalues Are Well Fit by the LML-RMT Predictions. In each panel, the grey histogram shows the bulk eigenvalue distribution of the scaled count matrix computed from the metacell model (top row) or the true scaled count matrix (bottom row) for four of the datasets. The curves show the eigenvalue distribution according to our LML-RMT predictions (solid blue) and the Marchenko-Pastur distribution (dashed red).

Across all the datasets, the MP distribution is a bad fit, seeming to contradict the results of Aparicio et al. However, Figure 3A shows the MP distribution against the true distribution as the number of genes varies through 1000, 2000, and 5000. As the number of genes increases, the MP fit improves. Aparicio et al. considered scaled count matrices with more than 10, 000 genes, much greater than the number of genes used by other groups in fitting the MP distribution. We can use the spike-bulk decomposition to understand this effect.

**Figure 3:**
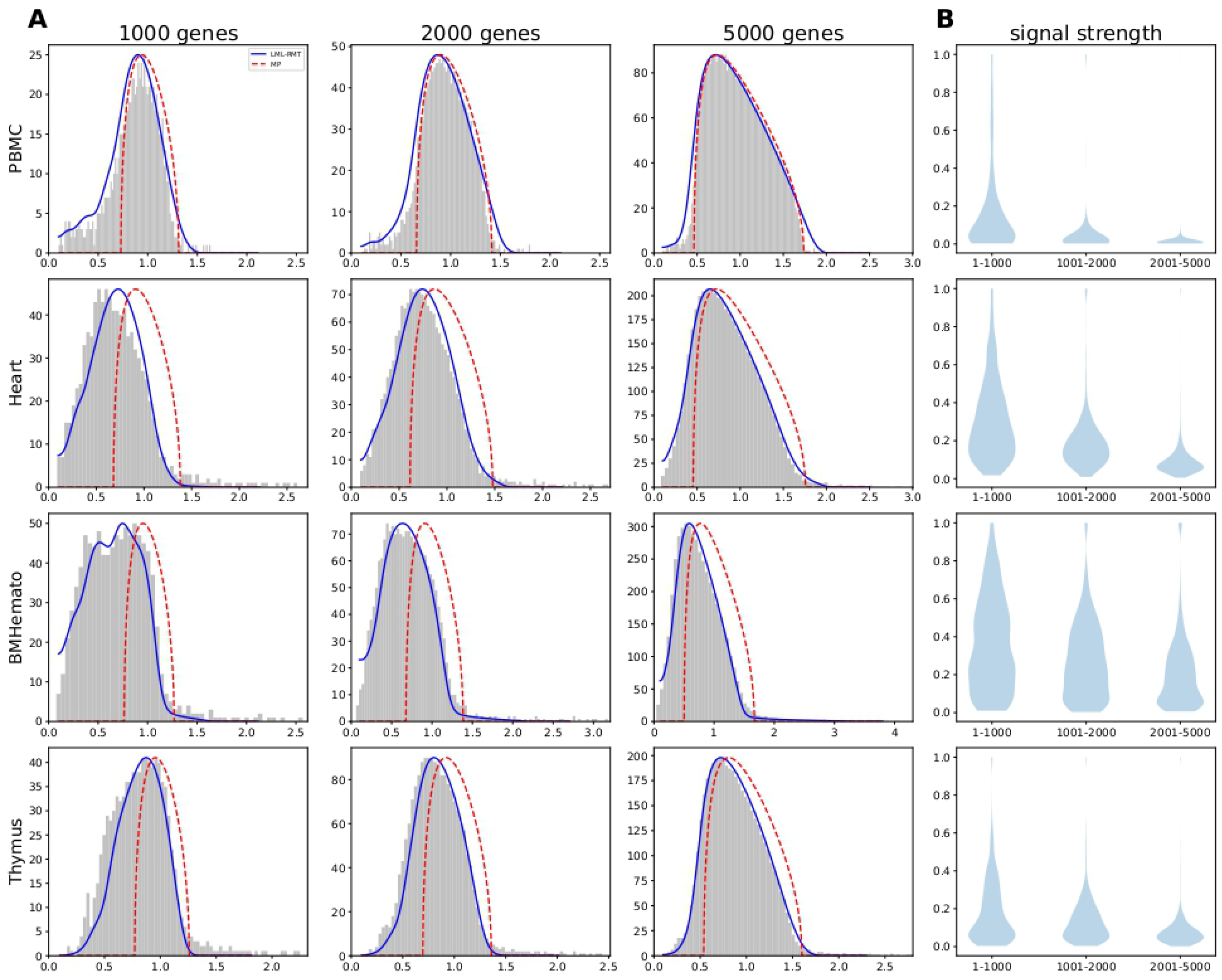
The Marchenko-Pastur Distribution Fit Improves as the Number of Genes Increases. (A) In each panel, the grey histogram shows the bulk eigenvalue distribution of the true count matrix and the eigenvalue distribution according to (blue curve) our LML-RMT predictions and (dashed red curve) the Marchenko-Pastur distribution. The columns show the number of genes included in the count matrix. (B) The distribution of the signal strength (see text for definition) for the genes of the 1000 gene count matrix (1-1000), for the genes of the 2000 gene count matrix not in the 1000 gene count matrix (1001-2000), and for the genes of the 5000 gene count matrix not in the 2000 gene count matrix (2001-5000).

Recall that the scaled count matrix is constructed by z-scoring the columns, making each column norm 1. If *S* = 0 and we assume iid normal entries of *B*, then the spectrum of *Z* will be MP distributed. But since *S* ≠ 0 for scaled count matrices, we cannot expect *Z* to be Marchenko-Pastur distributed. We can get an estimate for the column norms of *S* and *B* in the true dataset by using the *S* and *B* decomposition of *Z*_metacell_. Let *S*_*g*_ and *B*_*g*_ be the *g*th columns of *S* and *B*. Since each column of *Z* has norm 1, ∥*S*_*g*_∥^2^ + *E*[∥*B*_*g*_∥^2^] = 1, and we define the signal strength of gene *g* as ∥*S*_*g*_∥^2^. Figure 3B shows the distribution of the signal strength across the genes. In all the datasets shown, as the number of genes exceeds 1000, subsequent genes have much lower signal strength, meaning that the noise *B*_*g*_ captures most of the column’s norm, leading to a better fit of the MP distribution.

We next consider the outlier eigenvalues. To explain our approach, let *k* be the number of eigenvalues greater than the right bulk edge (i.e. the outlier eigenvalues) according to the LML-RMT predictions. We matched the *k* RMT predicted eigenvalues with the top *k* eigenvalues of 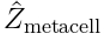and 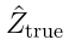. Figure 4 shows the empirical versus predicted values. As was the case with the bulk spectrum, the fit to the metacell eigenvalues is excellent (relative error 0.01). The error in the fit to the true eigenvalues is higher (relative error about 0.11).

**Figure 4:**
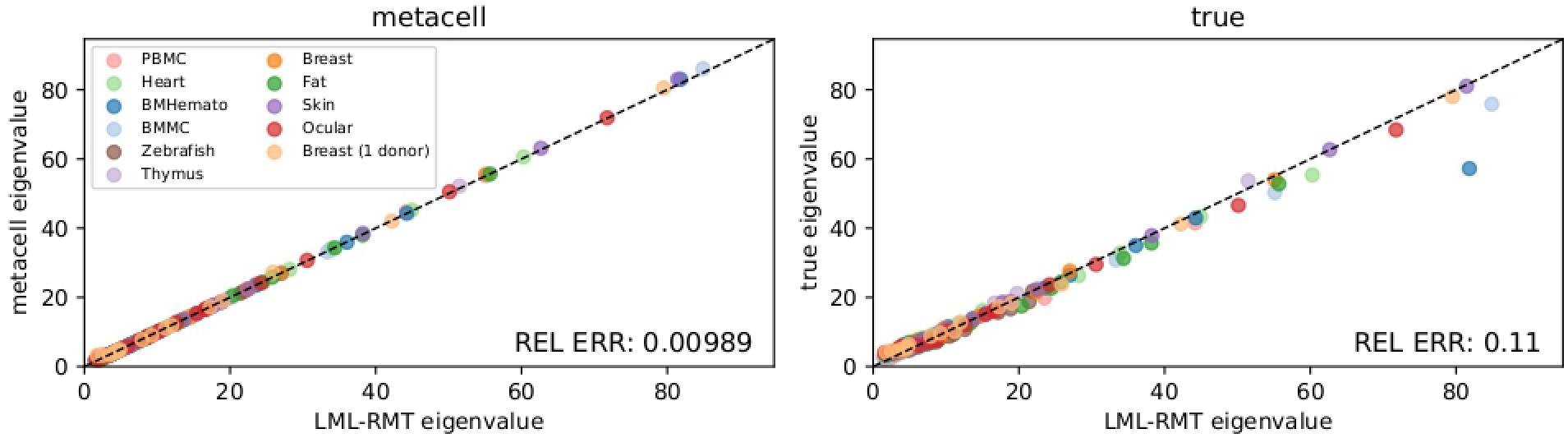
Outlier Eigenvalues are Well Approximated by LML-RMT Predictions. Each panel shows the predicted eigenvalue based on LML-RMT formulas (x-value) and the eigenvalue computed numerically for the particular scaled count matrix. All 11 datasets are included in each panel. REL ERR gives the average relative error of the prediction over all eigenvalues and datasets.

### 1.2 Geometry of the PCA Embedding Within Metacells

Now we turn to considering the PCA embedding. As noted above, under the LML-RMT prediction, the PCA embeddings of the cells in a given metacell are distributed as iid samples from an MVN. We start by investigating the fit of these predictions to individual metacells, below we consider the geometry of the metacells relative to each other. There are two related questions. First, for a particular metacell, are the PCA embeddings of the cells well fit by an MVN model? And second, if indeed metacells are well fit by an MVN, is the covariance structure of the metacell well fit by the LML-RMT predictions?

To address the first question, we test for normality using the Kolmogorov-Smirnov (KS) statistic. The KS statistic is a univariate statistic, comparing a univariate distribution to a univariate normal. For each metacell, we project the metacell’s cells onto 10 × *k* random unit directions in PCA space (recall *k* is the PCA space dimension) and test each projection for normality using the KS statistic. Since the KS distribution depends on the number of samples (here the number of cells in each metacell) and the large number of metacells and the PCA dimensionality raises multiple testing issues, we evaluated normality by comparing the KS distribution found for the PCA embedding generated by 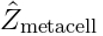 and by 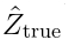 to the KS distribution generated by sampling directly from the MVN predicted by the LML-RMT. Figure 5 shows that the KS distributions for the true and metacell PCA embeddings largely overlap with those of the LML-RMT predictions.

**Figure 5:**
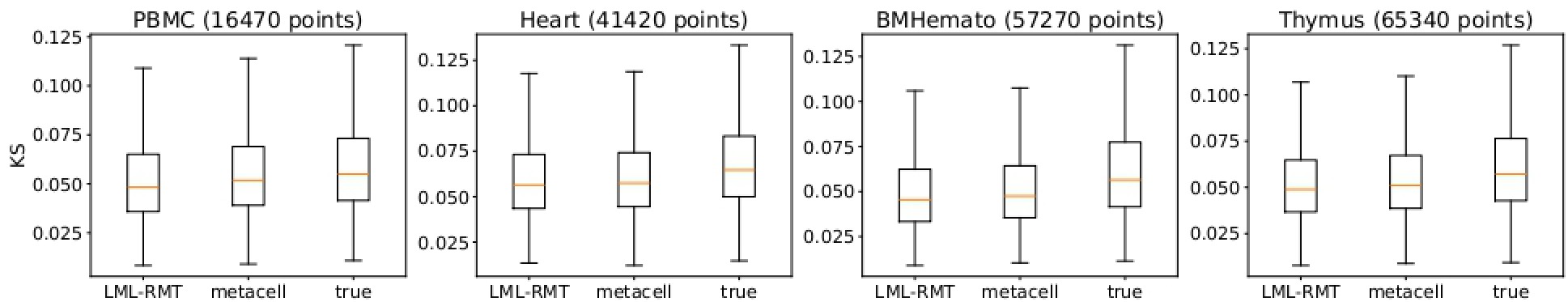
Kolmogorov-Smirnov Tests for the Normality of Metacell PCA Embeddings. Four of the datasets are shown. In each panel, the box plots give the Kolmogorov-Smirnov (KS) statistic over all metacells.

To address the second question, Figure 6 compares the variance of each metacell’s PCA embeddings formed from 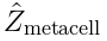 and 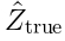to the LML-RMT predictions. Similar to the KS test, variances were computed by projecting cells onto different directions in PCA space. Also shown as a baseline is a comparison of the LML-RMT prediction against metacells formed from *Z*_normal_, which differ from the LML-RMT predictions only due to finite sample effects, see Methods for comments regarding the difference. We refer to results computed using *Z*_normal_ as results generated under the normal model. Since finite sample effects introduce stochasticity that is not our focus in assessing the LML-RMT predictions, we use the normal as a null against which to judge the relative error of the LML-RMT predictions. The fit for the metacell model is excellent and has the same relative error (about 0.09) as that for the PCA embedding of the normal model.

**Figure 6:**
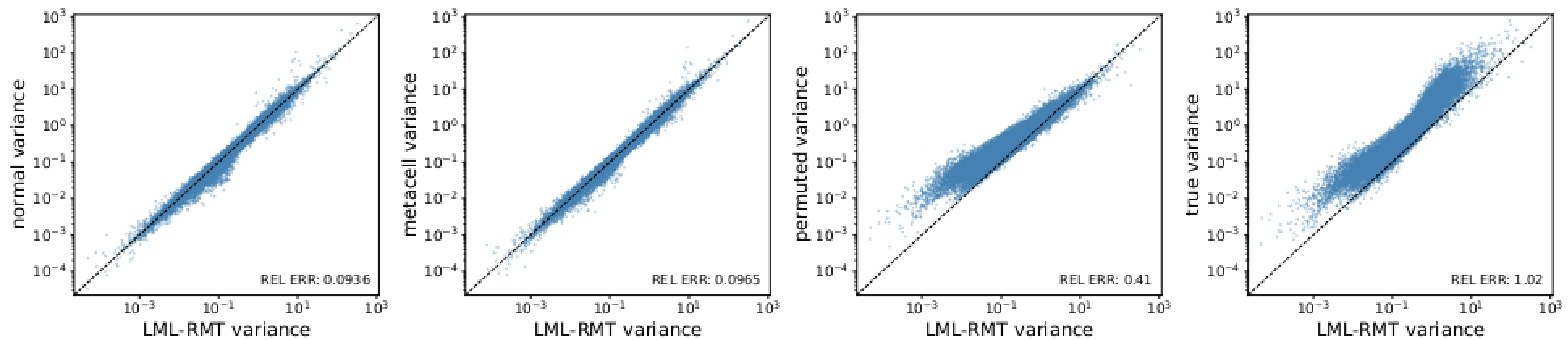
The Error in LML-RMT Predicted Variance for PCA Embeddings of Individual Metacells. Each panel compares the variance of cells within a metacell versus the LML-RMT predicted variance. REL ERR gives average relative error over datasets and metacells. Each point corresponds to a particular dataset, metacell, and direction. The variances produced for the normal model (far left panel) capture error only due to finite sample size effects in the LML-RMT predictions and serve as the null against which to judge the LML-RMT accuracy.

The LML-RMT predictions of the variance underestimate the true variances, with a relative error of about 1.0. There are two reasons the metacell model may fail to fit the true data. First, the negative binomial distribution might not fit the distribution of individual genes and, second, the true counts may have gene correlations within metacells. To tease out these effects, we defined a count matrix 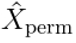by independently permuting columns (i.e. genes) of the true count matrix 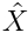within each metacell. 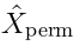has the same distribution of counts for each gene as 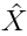, but the within metacell gene correlations are gone. We then applied the standard workflow to 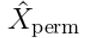 and repeated our analysis. As was the case for the true count matrix, the LML-RMT predictions underestimate the variance of the PCA embeddings for the permuted count matrix, although the underestimate is smaller, with a relative error of 0.41 versus 1.02 for the true data, suggesting that the metacells have genes that are not well fit by the negative binomial model, but that the majority of the error comes from the absence of gene correlations within metacells under the metacell model.

### 1.3 Geometry of the PCA Embedding Across Metacells

In most downstream applications, the PCA embedding’s geometry across metacells is more important than the geometry of individual metacells. For example, in the context of clustering, a metacell that is far away from all other metacells will form its own cluster, regardless of the variation and geometry of its embedding. Similarly, for batch correction, what matters is the relative geometry across batches of metacells that represent the same underlying cell states. To explore this issue, we consider a metacell and its closest metacell neighbors, where the means of the metacells in PCA space were used to assess distances between metacells. For the LML-RMT predictions, we used the analytic values of the means. For the normal, metacell, permuted and true PCA embeddings, we used the sample means. The overlap between metacells can be quantified by the distance between the metacell and its neighbors scaled by the standard deviation of the metacell along the line connecting the means. Figure 7 compares the predicted and true values of this ratio (see the figure for an exact formula of the ratio) for the 5 closest neighbors of each metacell.

**Figure 7:**
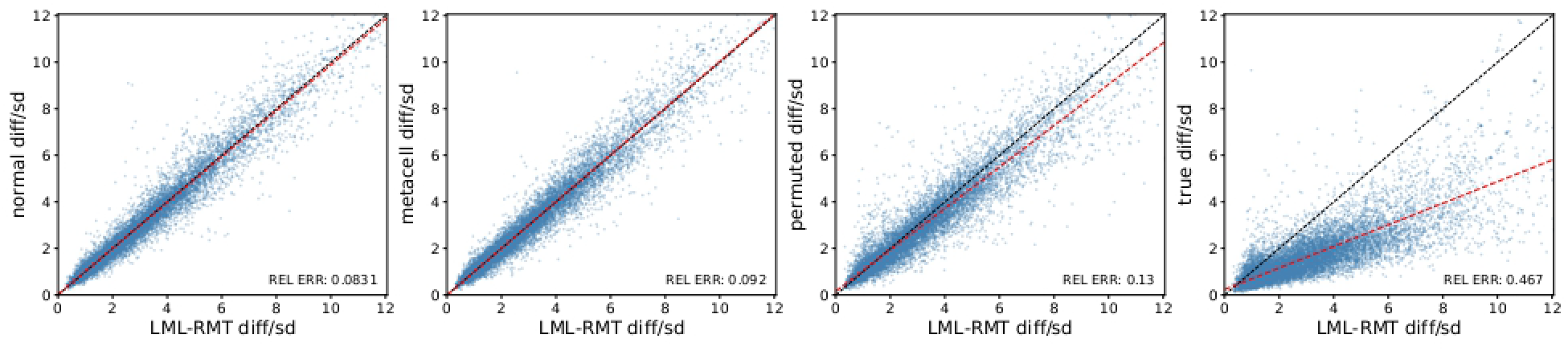
Scaled Distance from Metacell to Its Neighbors. Given a metacell and its neighbor with means *µ* and *µ*^*′*^ in PCA space, respectively, we compute the ratio∥ *µ* −*µ*^*′*^ ∥*/σ* where *σ* is the standard deviation of the metacell in the direction of *µ*− *µ*^*′*^. Each point in each panel shows the ratio for a metacell and one of its 5 closest neighbors according to the LML-RMT predictions (x-value) versus the ratio for the same metacell and neighbor based on the actual PCA embedding (y-value). The red dashed line shows the linear regression fit to the points.

The relative error of the LML-RMT predictions is roughly 0.09 for both the normal and metacell model PCA embeddings, showing that the LML-RMT predictions are accurate for the metacell model up to an error associated with finite sample size. The relative error for the permuted count matrix is 0.13 versus 0.47 for the true count matrix, showing again that gene correlation within the metacell is the main driver of error in the LML-RMT predictions for true datasets. But further, LML-RMT is roughly an unbiased estimator in the case of the permuted count matrix, but is upward biased for the true count matrix. We conclude that true gene counts that violate the negative binomial assumption of the metacell model lead to noise that is not localized in a particular direction, while noise due to gene correlations is correlated with the direction from a metacell to its closest neighbors.

We now turn to investigate the global geometry of the metacells. For each PCA embedding, we compute the k-nearest neighbor graph (knn) for the cells using 10 nearest neighbors. Then for each ordered pair of metacells, metacell *i* and metacell *j*, we compute the fraction of edges emanating from cells in metacell *i* that end in a cell in metacell *j*. We compute these fractions across all datasets and metacell pairs using the LML-RMT predictions and using the normal, metacell, permuted and true scaled count matrices. Figure 8 shows the fits of the LML-RMT predictions.

**Figure 8:**
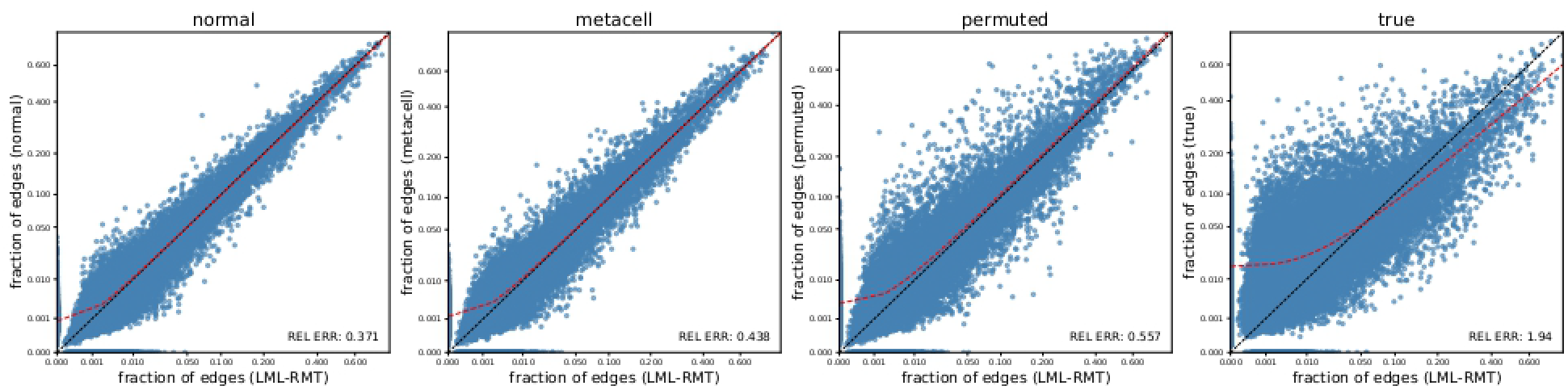
LML-RMT Predictions of the k-Nearest Neighbor Graph Across Meta-cells. For each PCA embedding, we constructed the k-nearest neighbor graph of the cells (10 nearest neighbors) and then computed for each ordered pair of metacells *i* and *j*, the fraction of edges emanating from cells in metacell *i* that end in a cell in metacell *j*. Each point in each panel shows this fraction for a metacell pair within a particular dataset according to the LML-RMT predictions (x-value) versus the fraction based on the PCA embedding of the noted count matrix (y-value).

As above, the relative error of the LML-RMT predictions is similar for the normal and metacell count matrices (0.37 and 0.44 respectively). LML-RMT predictions for the permuted count matrix have higher relative error (0.56) but are unbiased while predictions for the true count matrix have a much higher relative error (1.94) and were biased down, again reflecting predictions of metacells with more internal edges in the knn graph than the true metacells.

Here we see a small bias in the LML-RMT predictions relative to the metacell model values. Most of this bias is associated with small metacells, *<* 60 cells, which are connected to the other metacells in a stochastic way. This underlying stochasticity of the knn graph is seen in the relative error of 0.37 for the knn generated for the normal model. Table 2 shows the knn metric presented in Figure 8 split by dataset, with the datasets ordered from least to greatest relative error for the true count matrix. For the normal model, the table reports the mean relative error over 20 replicates, each formed from an independent sample of *Z*_normal_. The relative errors for the knn metric across these replicates vary as shown in the table. The metacell relative errors are within the range of the replicate relative errors for 4 of the 11 datasets and are slightly above the maximum replicate relative error for the others.

**Table 2:** Relative error of the LML-RMT predictions of the k-nearest neighbor graph across metacells (as described in Figure 8), by dataset. The normal column gives the mean of this error over 20 replicate draws of *Z*_normal_, each taken through the PCA and k-nearest neighbor steps in the same way, and the range in brackets is the smallest and largest value over the replicates. Rows are ordered by increasing error for the true column.

| Dataset | Cells | Donors | Cell types | Stage | normal | metacell | permuted | true |
| --- | --- | --- | --- | --- | --- | --- | --- | --- |
| PBMC | 39,074 | 1 | 10 | diff | 0.37 [0.36, 0.38] | 0.43 | 0.50 | 0.60 |
| Heart | 29,938 | 7 | 14 | diff | 0.46 [0.43, 0.50] | 0.50 | 0.56 | 0.82 |
| Breast (1 donor) | 29,737 | 1 | 15 | diff | 0.34 [0.31, 0.37] | 0.35 | 0.53 | 1.11 |
| Zebrafish | 33,614 | 31 | 10 | dev | 0.44 [0.40, 0.47] | 0.53 | 0.80 | 1.12 |
| Ocular | 29,547 | 1 | 12 | diff | 0.34 [0.32, 0.36] | 0.40 | 0.50 | 1.27 |
| Breast | 182,192 | 55 | 16 | diff | 0.38 [0.36, 0.40] | 0.46 | 0.50 | 1.50 |
| Skin | 35,075 | 1 | 15 | diff | 0.32 [0.30, 0.34] | 0.35 | 0.40 | 2.00 |
| BMHemato | 61,038 | 4 | 69 | dev | 0.36 [0.33, 0.39] | 0.40 | 0.50 | 2.37 |
| Thymus | 64,081 | 20 | 16 | dev | 0.49 [0.47, 0.50] | 0.52 | 0.61 | 2.74 |
| Fat | 19,330 | 1 | 14 | diff | 0.34 [0.29, 0.38] | 0.36 | 0.58 | 3.00 |
| BMMC | 2,517 | 1 | 9 | diff | 0.32 [0.20, 0.64] | 0.52 | 0.66 | 4.80 |

### 1.4 Accuracy of the Metacell Model

Our results show some variability in the fit of the metacell model to true datasets. Table 2 raises the question of whether datasets with particular properties are better fit by the metacell model, at least as measured by the accuracy of the LML-RMT predictions. We lack enough datasets to make statistically significant inferences, but Table 2 suggests two possible trends: the two datasets with the fewest cells have the largest error and the three datasets with the smallest error are differentiated. We speculate that cell types in differentiated datasets have less internal variation which allows for denser sampling. Increasing the number of cells would also lead to denser sampling, leading us to speculate that accuracy of the metacell model depends on sampling density. More work is required to investigate this hypothesis.

As datasets become bigger, they are generally split into more batches. But batches also introduce more noise into the dataset, so their effect on the accuracy of the metacell model is not intuitive. The Breast (1 donor) dataset is the subset of the Breast dataset formed by the cells of a single donor. Although it has roughly one sixth of the cells of the Breast dataset, its error is smaller, suggesting that variation across donors degrades the fit of the metacell model more than the reduced number of cells does. Importantly, we have not batch corrected our datasets. Batch correction might improve the accuracy of the metacell model. More work is required.

### 1.5 The Range of Metacell Variance Across a Dataset

Under the metacell model, any variance in the PCA embedding of cells within a metacell arises due to negative binomial noise and can be labeled as noise. We next investigate whether some metacells have more variance than others. Given a dataset, for each meta-cell we compute its cells’ variance in PCA space along the metacell’s direction of maximal variance. Since we are interested in the range of variance across the metacells composing a given dataset, we further normalize by scaling the smallest metacell variance to 1. Figure 9 shows that variance across metacells within a dataset ranges over more than 10 fold and a few metacells have variance more than 50 fold greater than the minimum. As seen in the figure, the range of variance seen under the metacell model and for true datasets is similar, suggesting that while the metacell model underestimates the true variance, it captures the range of within metacell variance. The range of metacell variance seen within datasets suggests that tools that depend on PCA distance may be confusing noise for signal and that a statistical model of PCA embeddings can lead to improved scRNAseq analyses.

**Figure 9:**
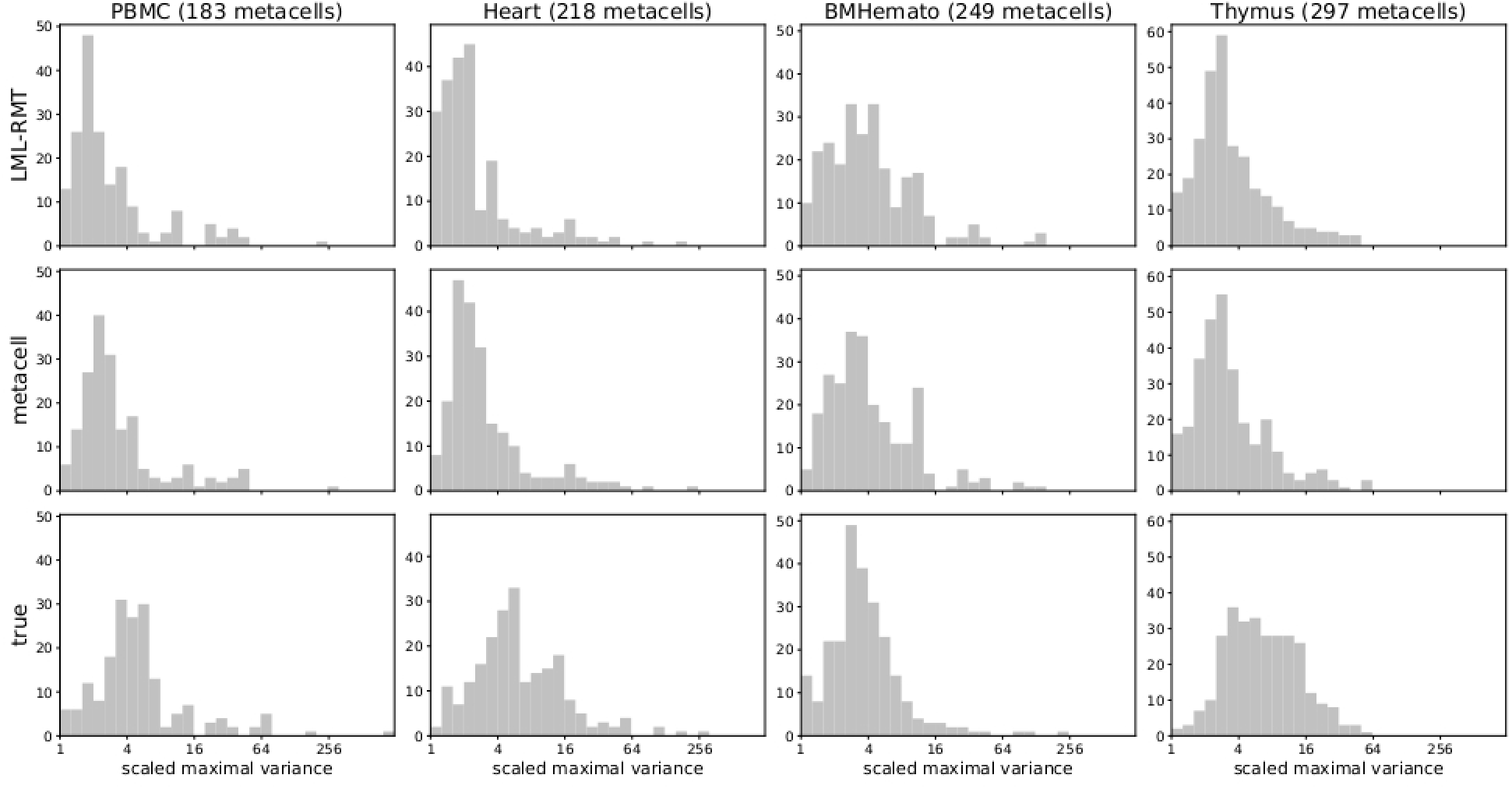
The Range of Metacell Variance in PCA Space Within Datasets. Panels show the scaled variance across all metacells with columns specifying the dataset and rows showing the scaled variance for the LML-RMT prediction (top) metacell model (middle) and true counts (bottom).

## 2 Discussion

In this work, we have used RMT tools to connect a statistical model of scRNAseq cell counts to a statistical model of cell embeddings formed by a common scRNAseq workflow - what we term the standard workflow - that log-normalizes, z-scores and then embeds cells using a PCA. We have shown that under a metacell model of cell counts, the standard workflow leads to cells distributed as a GMM in PCA space. We have validated this result by fitting the metacell model to 11 datasets of various form and size, generating count data according to the metacell model, and then showing that the GMM predictions fit the PCA embedding of the generated counts well.

In the process of our analysis of the metacell model, we have established novel results at the intersection of RMT and scRNAseq. We have shown that the eigenvalues of a scaled count matrix generated by negative binomial count noise in the parameter regime of scRNAseq datasets are well modeled by eigenvalues generated from Gaussian noise. We have resolved a seeming contradiction in the RMT-scRNAseq literature by showing that the fit of the Marchenko-Pastur distribution to the bulk spectrum of scaled count matrices improves as the number of highly variable genes included increases, thereby explaining why some groups have reported a Marchenko-Pastur distribution for scRNAseq datasets while others have reported deviations. We have provided the first spike-bulk model of scRNAseq datasets and we have extended RMT tools to account for the correlation between the spike and bulk that exists in our model.

The characterization of the metacell model’s cell embeddings as GMM distributed provides a framework through which to validate the metacell model itself, at least in terms of its PCA embeddings. We show that the metacell model leads to underestimates in the variance of true cell embeddings and through numerical experiments we show this underestimate arises due to gene correlation within the metacells.

Intriguingly, we noted better prediction of true cell distributions for differentiated datasets as compared to developmental datasets and poorer prediction for datasets with small sample size. We speculate that the scRNAseq sampling of differentiated datasets is denser because there is less variation in cell expression, which makes the metacell model more accurate. Raising the number of cells sampled would similarly increase density, but further work is needed to explore this issue. Importantly, the one dataset that fit the metacell model best is the Zheng et al. PBMC dataset, a well-known dataset used in many benchmarking studies. Our results suggest that this dataset is not typical and extrapolation of results from it should be viewed with caution.

While the metacell model underestimates the absolute variance of true metacells in PCA space, it captures the relative variance across metacells. We find that metacell model generated and true PCA embeddings have metacells that range in their internal variance by similar factors with the majority of the metacells covering a 10-fold range, while some outlier metacells have more than 50-fold variance relative to the metacell with minimal variance. Downstream tools that treat distances in PCA space uniformly, such as clustering and nearest neighbor graph construction, may therefore mistake the noise of high variance metacells for biological signal.

Previous authors have constructed distributions for cell embeddings. Variational autoen-coder models such as scVI [40, 19] assume a prior on their embedding spaces and, after fitting to a dataset, a posterior for the embedding space can be computed. However, these distributions do not have an explicit connection between a count model and the embedding distribution so the posterior does not reflect noise in the dataset itself. While scVI uses the negative binomial as part of its generative model, the degree to which this captures different aspects of noise in a dataset is unclear. For example, if we sample a dataset from a particular count model twice and apply scVI to each, the posterior will be different for the two generated cases. Putting aside the analysis of other embeddings, we are not aware of any previous analysis of cell distribution arising from the standard workflow, despite its common usage.

Analysis of PCA embeddings connects naturally to RMT, which mostly focuses on the eigenvalues of random matrices. Previous groups have empirically investigated the eigenvalue distribution of scaled count matrices. Several groups have developed methods based on RMT analysis of spike-bulk models [46, 33, 54, 15, 29, 32], but these methods have not modeled the spike and have assumed RMT results that depend on isotropy of the bulk relative to the spike and the Gaussian universality of the bulk matrix. For our metacell model analysis, the bulk is not isotropic relative to the spike, a feature that is likely true for scRNAseq datasets in general. The application of RMT methods to specific count matrix models provides the potential to use RMT to characterize scRNAseq embedding spaces, with significant implications for improvement of scRNAseq algorithms.

There are several limitations of our work. Perhaps the most significant is that there is no current algorithm for constructing metacells. We have used existing algorithms as a basis for our metacell construction, but we cannot rule out the possibility that there is some choice of metacells that leads to a better fit of the metacell model to true datasets. In another direction, we have not batch corrected our datasets. How batch correction affects the accuracy of the metacell model as a predictor of PCA embeddings for true datasets is unclear. More datasets are needed to explore these questions in a rigorous manner. In terms of the LML-RMT analysis, we have not explored the role of library size in the fit. Our metacell model assumes a fixed library size and the contribution of this assumption to the metacell model fit of true data is unclear.

## 3 Methods

All code used to generate results, figures, and tables presented in this manuscript is available at https://github.com/SLeviyang/metacellGMM

### 3.1 Datasets

We downloaded a count matrix for each dataset. In some cases we selected a subset of the dataset’s cells. In all cases, we applied a QC step that filtered cells for UMI count and mitochondrial gene expression and filtered genes for total counts. Table 3 provides download, subsetting and QC information for each dataset. For each dataset, we then selected the 1000, 2000 or 5000 most highly variable genes using scanpy’s `highly_variable_genes` function with the `seurat_v3` flavor. We used scanpy to implement log-normalization and z-scoring through the following code,

**Table 3:** Data sources and cell filtering. Source is the CELLxGENE dataset identifier or the repository accession of the downloaded counts. QC cutoffs are the thresholds applied to the raw counts: the range of total UMI counts and of genes detected per cell, the maximum percentage of mitochondrial (MT) reads, and the gene filter. The last column describes any selection of cells made before those cutoffs were applied.

| Dataset | Source | QC cutoffs | Cell selection before QC |
| --- | --- | --- | --- |
| PBMC | 10x Genomics Fresh 68k PBMCs (Donor A), Zheng et al. 2017 | MT < 5%; UMIs 1,000–3,000; genes with > 50 counts. | None; all 68,579 cells. |
| Heart | CELLxGENE 49312f8c-a75a-4672-9c8a-9078acd35bdb (Cells of the adult human heart) | Authors' QC only; genes with > 50 counts. | Single-cell libraries only: cells with `cell_source =` Sanger-Cells (45,885 of 486,134); nuclei and CD45<sup>+</sup>-enriched libraries excluded. |
| BMHemato | GEO GSE245108 | UMIs 1,000–75,000; genes 500–8,000; MT ≤ 15%; genes with ≥ 50 counts. | RNA counts of the 71,994 annotated cells in the 12 GEO filtered matrices; ADT features dropped. |
| BMMC | GEO GSE194122 (NeurIPS 2021 multiome BMMC) | UMIs 500–30,000; genes 200–8,000; MT ≤ 20%; genes with ≥ 50 counts. | Batch s1d1 only, GEX features only; cells with ≥ 100 genes and genes in ≥ 100 cells (6,224 cells). |
| Zebrafish | Zebrahub, figshare DOI 10.6084/m9.figshare.20510367 | UMIs 1,000–20,000; genes 500–6,000; MT ≤ 10%; genes with ≥ 50 counts. | None; all 120,444 cells (bud stage to 10 dpf). |
| Thymus | Zenodo DOI 10.5281/zenodo.3572421 (Park et al. 2020) | UMIs 2,000–30,000; genes 800–5,000; MT ≤ 6%; genes with ≥ 50 counts. | The 76,994 cells of the authors' annotated T cell/thymocyte subset, with raw counts taken from the full-atlas count matrix. |
| Breast | CELLxGENE 2df9adc8-fefc-4a96-9d80-0a7c6b4576e1 (HBCA global) | UMIs 800–50,000; genes 500–6,500; MT ≤ 8%; genes detected in ≥ 100 cells. | None; after QC, 200,000 of the 782,667 passing cells drawn at random. |
| Fat | CELLxGENE 5e5e7a2f-8f1c-42ac-90dc-b4f80f38e84c (Tabula Sapiens, fat) | UMIs 2,000–50,000; genes 500–8,000; MT ≤ 20%; genes with ≥ 50 counts. | Donor TSP25 only, brown adipose tissue libraries only (6 lanes, 40,619 cells). |
| Skin | CELLxGENE f0f0d7c4-3bec-428e-9539-c99d36548d96 (Reynolds et al. 2021) | UMIs 1,000–30,000; genes 500–5,000; MT ≤ 10%; genes with ≥ 50 counts. | Donor S4 only (50,492 cells). |
| Ocular | CELLxGENE 8560f809-5794-484e-9152-b63efa91d134 (ocular anterior segment, scRNA-seq) | UMIs 600–40,000; genes 400–6,000; MT ≤ 5%; genes with ≥ 50 counts. | Donor BCM_23_0131 only (42,612 cells). |
| Breast (1 donor) | CELLxGENE 2df9adc8-fefc-4a96-9d80-0a7c6b4576e1 (HBCA global) | UMIs 800–50,000; genes 500–6,500; MT ≤ 8%; genes detected in ≥ 100 cells. | Donor HBCA_Donor_33 only (39,446 cells); no downsampling. |

~~~
sc.pp.normalize_total(s, target_sum=1e4)
sc.pp.log1p(s)
sc.pp.scale(s)
~~~

where s is the scanpy object for the dataset and sc is the scanpy package. We then used numpy’s svd function to compute the PCA embedding.

### 3.2 Metacell Construction

Given a dataset, we partitioned cells into metacells in a two step process. In forming the metacell partitioning we exclusively used the true counts of the dataset. First, we used Supercell [13] applied to the counts of the highly variable genes to form an initial partition. The granularity of Supercell is set by its graining level *γ*, the average number of cells per supercell. Rather than fix *γ* directly, we fixed a target number of supercells *m*_0_ for each dataset and set *γ* = *C/m*_0_, where *C* is the number of cells in the dataset. We used *m*_0_ = 80 for most datasets, *m*_0_ = 120 for BMHemato, Thymus and Breast, and *m*_0_ = 500 for Zebrafish. We ran Supercell separately within each annotated cell type, so that every supercell contains cells of a single cell type, and apportioned *m*_0_ across the cell types in proportion to their cell counts. Cell types with fewer than 100 cells were excluded. Supercell was run with its defaults of 5 nearest neighbors and 1000 variable genes. Supercells with fewer than 100 cells were discarded.

We noted that the metacells produced by Supercell did not have an MVN distribution in PCA space. Since the metacell model implies that the PCA embedding of each metacell is an MVN, we included a second step in which we refined the Supercell partition by splitting supercells with non-normal PCA embeddings. This step was included to potentially optimize the fit of the metacell model PCA embeddings to true embeddings. To be sure that the accuracy of our LML-RMT predictions was not associated with a particular metacell construction, we also constructed a metacell partition that did not include the second partition step and was composed of very coarse metacells and then repeated the analysis presented in the Results. See the bottom of this section for details.

In the second step of our two-step metacell construction, the refinement proceeds in rounds. In each round, we compute a Kolmogorov-Smirnov (KS) statistic for each metacell by projecting its cells onto the principal directions of its sample covariance matrix in PCA space, computing the KS statistic of each projection as in the Results, and taking the max over the *k* directions. We z-score the statistic using its mean and standard deviation under the null of normality for that metacell. This gives us a z-scored KS statistic for each metacell. In each round, every metacell with a z-score greater than 3 and a KS statistic greater than 0.05 is split in two along the principal direction with the largest KS statistic. (The second condition prevents the splitting of large metacells whose departure from normality is statistically significant but negligible in size.) We fit a two component Gaussian mixture to the projections of its cells onto that direction and assign each cell to one of the two children by sampling from its posterior membership probabilities. The split is accepted if at least one child has at least *n*_min_ = max(40, *k* + 20) cells, where *k* is the PCA dimension, and a *z*-score below that of the parent; otherwise the parent is marked as refractory and is removed from the dataset. Metacells with fewer than *n*_min_ cells are also removed. Rounds continue until no metacell is flagged or no split is accepted, up to a maximum of 20 rounds.

All of our datasets contained cell type labels. As a comparison to the 2-step metacell partition, we also partitioned the cells by cell type so that each metacell corresponded to a single cell type. Results were qualitatively similar for this partitioning, except that the true metacells were not roughly normally distributed in PCA space. Figures S1-S5 of the Supplementary Information repeat Figures 4-8 for this partitioning. Importantly, using this coarse metacell construction, we still found that the metacell model’s PCA embeddings were well described by the LML-RMT results, showing that our description of the metacell model as GMM distributed in PCA space is not dependent on a particular metacell construction.

### 3.3 Computing the spike-bulk decomposition

From equation (4), the distribution of *X*_*ig*_ is known. From equation (1), we numerically compute *E*[*Y*_*ig*_] and *V* [*Y*_*ig*_] for every entry of *Y* (here *V* [] is the variance). Then to describe the distribution of *Z* using (2), we compute,

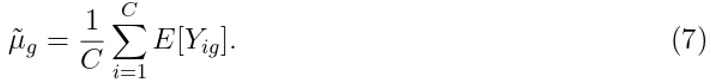

Note that 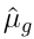 in (2) is formed as the sample mean in the standard workflow, while 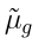 is the true mean. Similarly, we compute

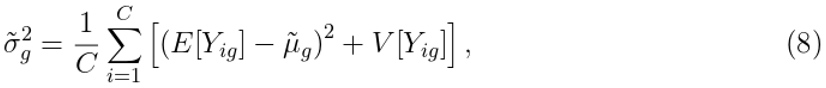

where here again the *V* [*Y*_*ig*_] terms make 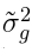 a true variance instead of a sample variance. Finally, we form the decomposition *Z* = *S* + *B* using

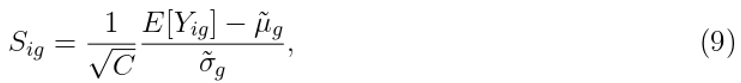

and

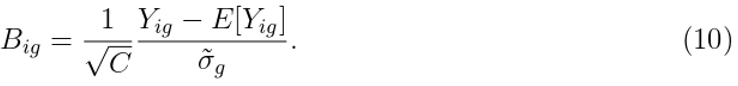

*B* is mean-zero by construction and since we know *V* [*Y*_*ig*_], we can compute *V* [*B*_*ig*_]. In the large sample limit, 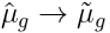 and 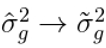, so our LML-RMT predictions are not affected by these approximations and the error arising from embeddings based on *Z*_normal_ is associated only with finite sample effects.

Two properties of *S* will be important in the development of the LML-RMT predictions below. First, notice that the columns of *S* are centered. Second, since rows within a metacell have the same distribution, for any column of *S* the entries within a given metacell are identical.

### 3.4 Computing the LML-RMT Predictions

Given *S* and *B* we can form *B*_normal_ and (6) defines the spike-bulk decomposition for *Z*_normal_. Here, we describe our analysis and corresponding computations for the singular values of the bulk matrix *B*_normal_ and the outlier singular values and vectors of *Z*_normal_.

We use the results of Benaych-Georges and Nadakuditi and Benaych-Georges and Couillet, hereafter referred to as BG-N and BG-C respectively. Since the bulk *B*_normal_ is not isotropic relative to *S*, we need to extend the results of BG-N and we do so by exploiting results developed in BG-C. Using the notation of BG-C, two matrices play a key role in the analysis,

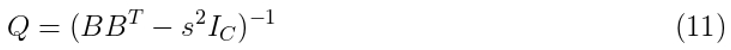

And

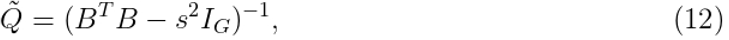

where *Q*, 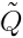match the notation of BG-C (BG-C have their samples in the columns while ours are in the rows, so their matrices are transposed relative to ours). Below, the analysis involves computations of various quadratic forms involving *Q*, 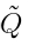which we compute using

Propositions 1.5 and 1.6 in BG-C. These two propositions describe results in a large matrix limit in which we take *G, C*→ ∞ with *G/C* fixed. Below all limits refer to this large matrix limit.

#### 3.4.1 Computing the Bulk Spectrum

The bulk matrix *B*_normal_ satisfies the assumptions of BG-C. Their Proposition 1.3 provides an iteration for computing the Stieltjes transform of *B*^*T*^ *B*, although we note that the seminal results of [60, 18] in the RMT analysis of mixture models derived a similar iteration. We use the MIXANDMIX algorithm [17] to compute the eigenvalues of *B*^*T*^ *B*. The algorithm computes the Stieltjes transform using the iteration, but with application of Anderson mixing to accelerate convergence, see [17] for details. Once the Stieltjes transform is computed, we compute the eigenvalue density by numerically inverting the Stieltjes transform.

#### 3.4.2 Computing the Outlier Eigenvalues

To compute the outlier eigenvalues, we first compute the SVD of 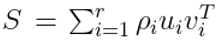, where *r* is the rank of *S*. We now follow a development similar to BG-N. Here we highlight key points and where our analysis differs, see their manuscript for full details. Define the (*C* + *G*) ×(*C* + *G*) matrix,

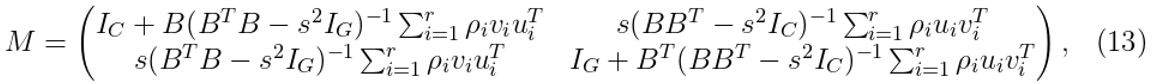

where *I*_*C*_ is the *C* ×*C* identity matrix and similarly for *I*_*G*_. Then *s* is a singular value of the scaled count matrix *Z*_normal_ if det(*M*) = 0 (here det is the determinant). Equivalently, by changing from the standard basis to that formed by the *u*_*i*_ and *v*_*i*_, we can construct a 2*r*× 2*r* matrix *M*^*′*^ with *s* solving det(*M*^*′*^) = 0. To define *M*^*′*^, we’ll take *r* = 2 to simplify notation; the case of general *r* is the obvious generalization. For the *r* = 2 case,

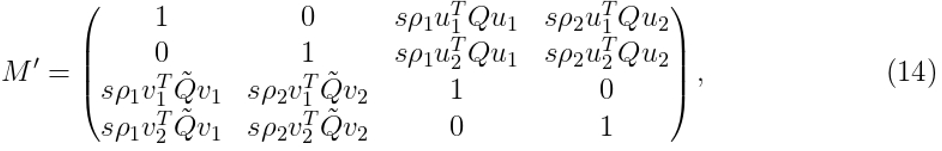

*M*^*′*^ is the analogue of the matrix in BG-N’s Lemma 4.1 with one important difference. The form of *M*^*′*^ depends on the large matrix limit 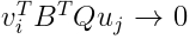 which BG-N justify and that also applies in our case. However, in the analysis of BG-N, the assumption that *B* is isotropic relative to *S* leads to 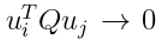 and 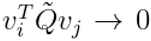for *i ≠ j* (recall → is the large matrix limit throughout) which does not hold in our case.

In BG-N, the isotropy assumption simplifies the structure of *M*^*′*^, leading to an explicit formula for the singular values (see BG-N’s Theorem 2.9). In our case, there is no such simplification. To find *s* we use a grid search. In practice, we compute the eigen-values rather than singular values using the scipy root finder within a grid search over [*b*^+^ + 0.1, max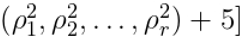 where *b*^+^ is the right edge of the bulk eigenvalues (recall *ρ*_*i*_ is the *i*th singular value of *S*). The rank of our spike *S* is the number of metacells minus one, which can be relatively large as seen in Table 1. However, since the metacells often share similar means with other metacells, most of the singular values of *S* are small relative to *b*^+^ and the number of outlier singular values ranges from 9 to 35, as shown in the PCA column of Table 1.

#### 3.4.3 Computing the Outlier Eigenvectors

For each outlier singular value *s*, we compute the left and right singular vectors of *Z*: *x* ∈ ℝ^*C*^ and *y* ∈ ℝ^*G*^ respectively. Again following BG-N, we decompose *x* and *y*,

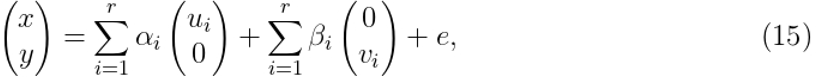

where *e* is orthogonal to the singular vectors of the spike. A simple computation transforms 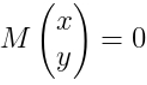 into

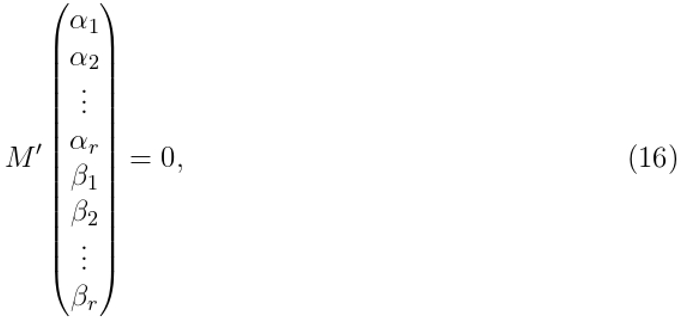

which gives the *α*_*i*_ and *β*_*i*_ up to a scalar. Using 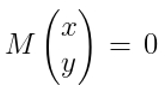 and pulling out the identity matrix in *M* gives

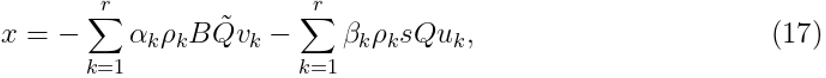

and the requirement ∥*x* ∥ ^2^ = 1 determines the scalar. As was the case above for the singular values, in BG-N these equations decouple, leading to explicit formulas for the *α*_*i*_ and *β*_*i*_ (see their Theorem 2.10) but in our case the more complex structure of *M*^*′*^ requires a numerical solution.

Specification of *x* and *y* still requires a description of *e*. We focus on *x* since *sx* forms a column of the PCA embedding matrix *W*, which is our object of interest. In what follows, we will need the following two properties of the *u*_*i*_. Let ***1*** be the *C* dimensional vector of all 1’s. Then for *i* = 1, 2, …, *r*

1. Each *u*_*i*_ is centered
2. The coordinates of *u*_*i*_ are constant within a metacell.

The first property follows from the centering of the columns of S, making *S*^*T*^ ***1*** = 0 and then plugging the SVD into ∥*S*^*T*^ ***1*** ∥ ^2^ gives the result. To see the second property, let *ϕ*_*a*_ be a vector that sums to 0 across indices within metacell *a* and is 0 for indices outside metacell *a*. Then since *S* has constant columns within a metacell, *S*^*T*^ *ϕ*_*a*_ = 0. Plugging in the SVD for *S* into ∥*S*^*T*^ *ϕ*_*a*_∥^2^ shows that 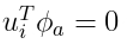 for all *i*, which gives the result.

We expand *e*,

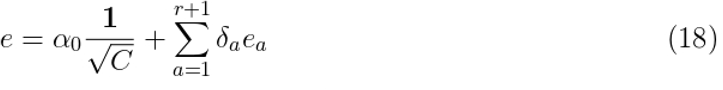

where *e*_*a*_ is a unit vector that is 0 for rows outside metacell *a* (recall there are *r* +1 metacells) and satisfies ***1***^*T*^ *e*_*a*_ = 0 and ∥*e*_*a*_∥ = 1. Applying this expansion transforms (15) into,

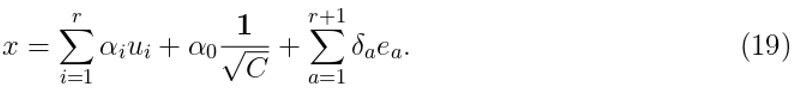

Using (17), we can compute *α*_0_:

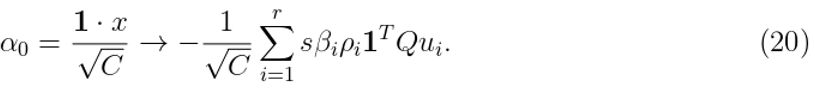

In our computations, we find that *α*_0_ is negligible and in practice can be ignored.

We now claim that each *e*_*a*_ is uniformly distributed on a unit sphere, i.e. Haar distributed. Let *C*_*a*_ be the number of rows in metacell *a* and define *O*_*a*_ as a rotation on 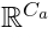with *O*_*a*_***1***_*a*_ = ***1***_*a*_ (here ***1***_*a*_ is the vector of all 1’s in 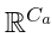). Define *O* as the extension of *O*_*a*_ to ℝ ^*C*^ that is the identity for rows outside metacell *a*. Since the columns of *S* are constant within a metacell and the columns of *B*_normal_ have constant variance within a metacell, *OS* = *S* and *OB*_normal_ ~*B*_normal_. This means *OZ*_normal_ ~*Z*_normal_ which implies *Ox*~ *x*. Since the *u*_*i*_ are constant within metacells, *Ou*_*i*_ = *u*_*i*_ and we find *O*_*a*_*e*_*a*_~ *e*_*a*_, which gives the Haar distribution.

Importantly, *e*_*a*_ is uniform not on the *C*_*a*_ dimensional unit sphere but on a *C*_*a*_ − 1 dimensional sphere, because of the restriction 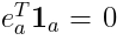. We simulate *e*_*a*_ as iid *N* (0, 1*/C*_*a*_), since for any fixed set of cells the entries 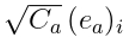 converge jointly in distribution to iid *N* (0, 1) in the large matrix limit.

To compute the large matrix limit of 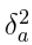, following the notation in BG-C, let *D*_*a*_ be the diagonal matrix in which all diagonal entries are 0 except for those corresponding to rows in metacell *a*, which have the value 1. Then,

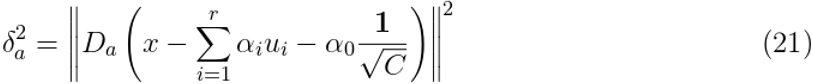

After expanding the right hand side of the equality above, the large matrix limit of each term can be computed using Propositions 1.5 and 1.6 of BG-C.

So far we have been dealing with a single singular value *s* and left singular vector *x*. To understand the distribution of a metacell, we need all the outlier singular values *s*^(1)^, *s*^(2)^, …, *s*^(*k*)^ and their associated left singular vectors *x*^(1)^, *x*^(2)^, …, *x*^(*k*)^. Using the superscript to denote the outlier index and letting *w*_*a*_ ∈ ℝ^*k*^ be the PCA embedding of a cell in metacell *a*, the above analysis gives,

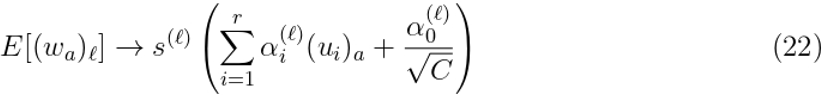

where by (*u*^(*l*)^)_*a*_ we mean the value of *u*^(*l*)^ in the *a*-th metacell, which is constant over the cells by the two properties of the *u*_*i*_ established above.

We also know the variance of the coordinates of *w*_*a*_,

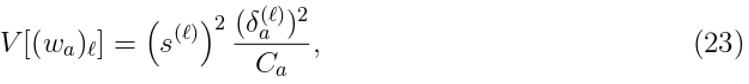

but we have left to compute the covariances across the PCA coordinates,

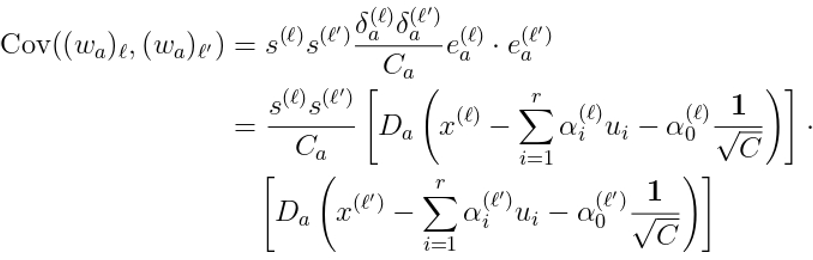

As in computing 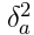, after substituting (17) for the *x* and multiplying the bracketed terms to the right of the last equality, the large matrix limits of all the terms can be computed using Propositions 1.5 and 1.6 of BG-C.

## 4 Supplementary Information

The figures of this section repeat Figures 4-8 of the main text with metacells defined by cell type rather than by the two step construction of the Methods. The LML-RMT predictions fit the metacell model as well as they do under the two step construction, while the true metacells, which are now whole cell types, depart further from the predictions.

**Figure S1:**
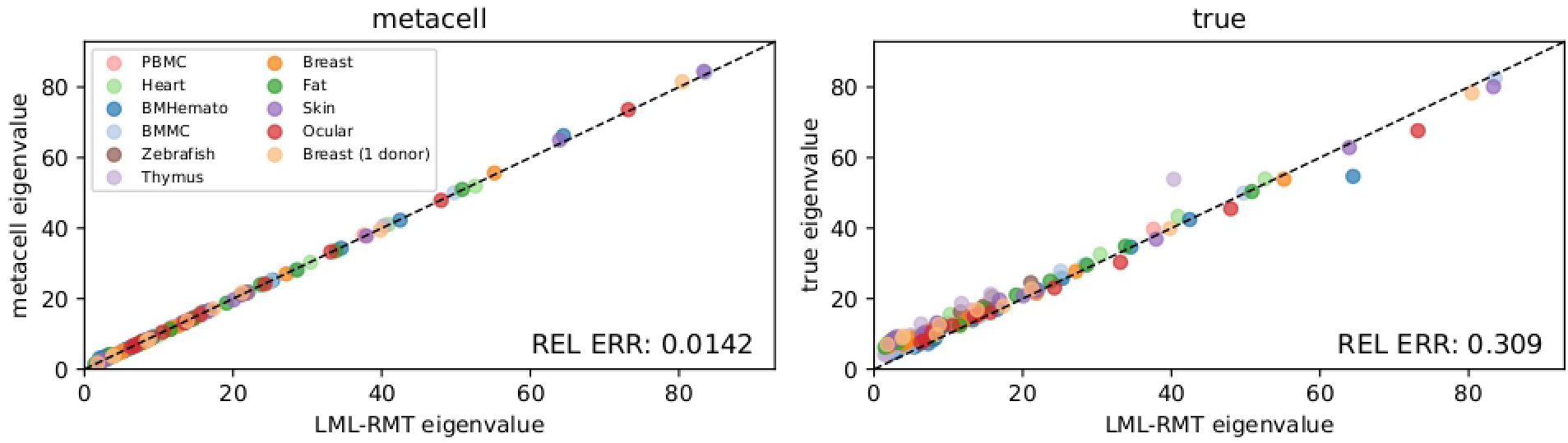
Outlier Eigenvalues for the Cell Type Partitioning. As Figure 4, but with each metacell formed by all cells of a given cell type.

**Figure S2:**
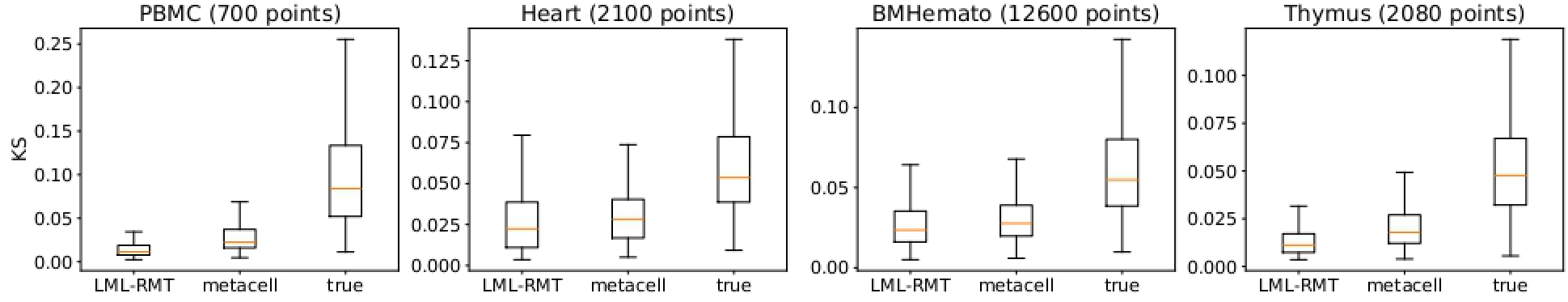
Kolmogorov-Smirnov Tests for the Normality of Metacell PCA Embeddings for the Cell Type Partitioning. As Figure 5, but with each metacell formed by all cells of a given cell type.

**Figure S3:**
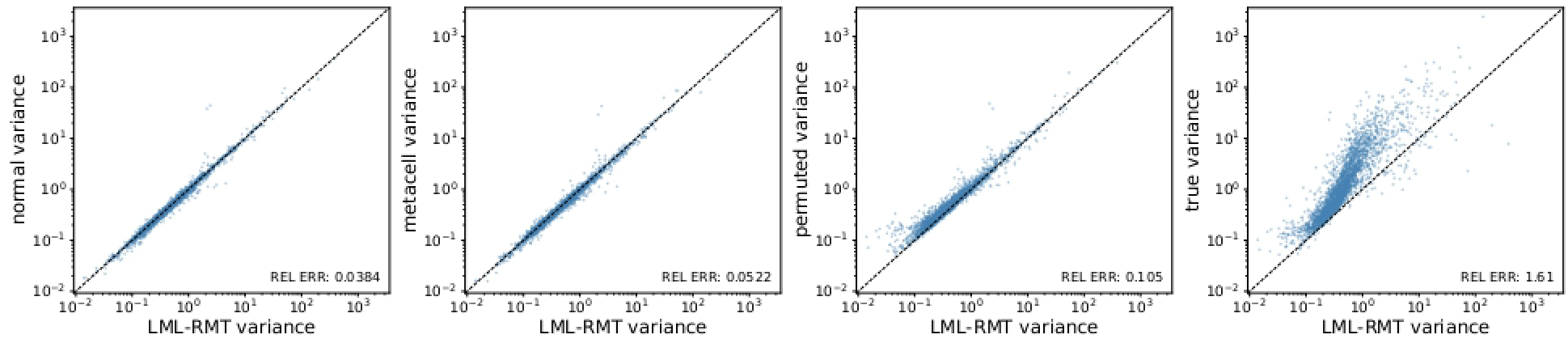
The Error in LML-RMT Predicted Variance for the Cell Type Partitioning. As Figure 6, but with each metacell formed by all cells of a given cell type.

**Figure S4:**
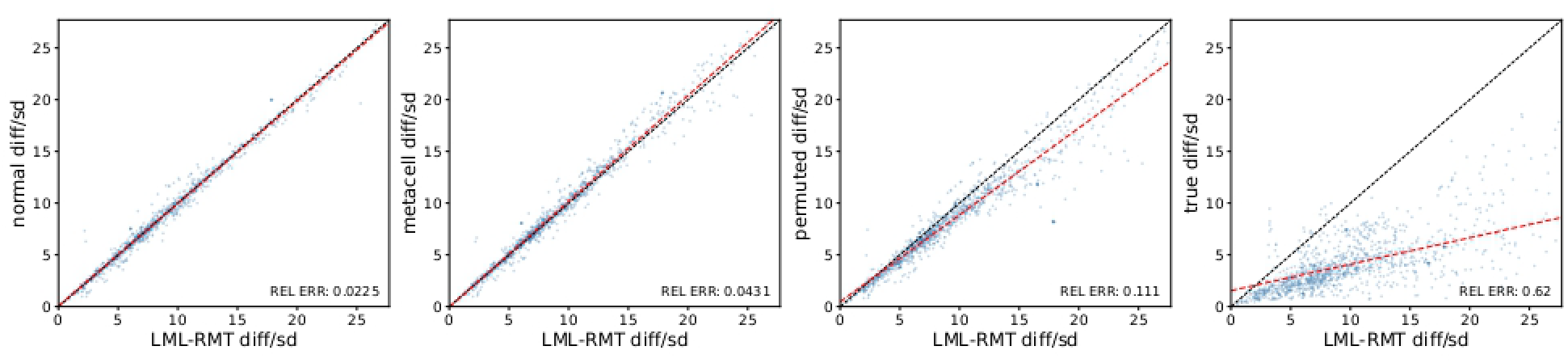
Scaled Distance from Metacell to Its Neighbors for the Cell Type Partitioning. As Figure 7, but with each metacell formed by all cells of a given cell type.

**Figure S5:**
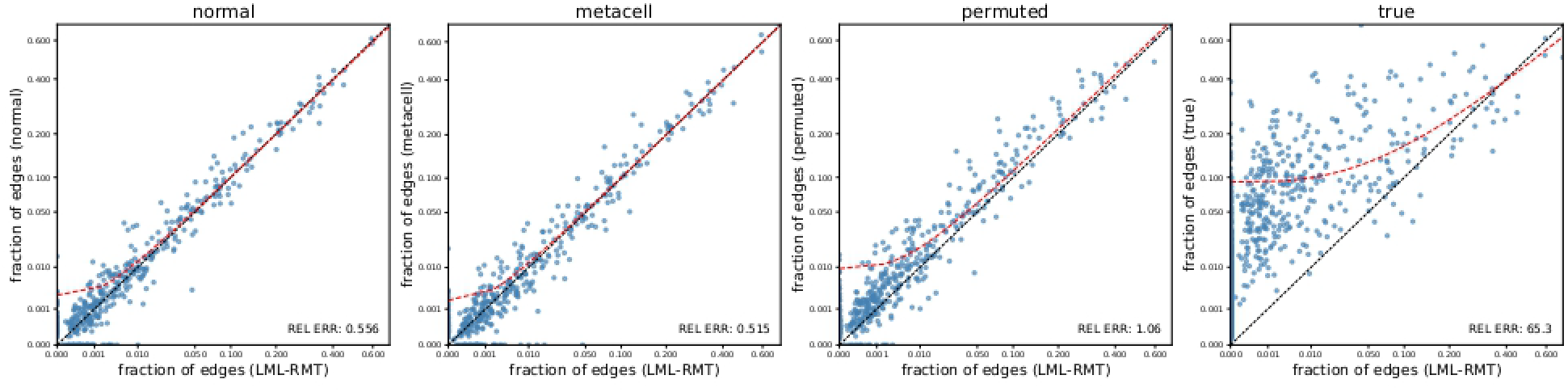
LML-RMT Predictions of the k-Nearest Neighbor Graph for the Cell Type Partitioning. As Figure 8, but with each metacell formed by all cells of a given cell type.

